# Multiplex-GCN Predicts Clinically Actionable Tumor Microenvironment Subtypes in Pancreatic Cancer

**DOI:** 10.64898/2026.10.22.761275

**Authors:** Shayori Bose, Paramita Basak Upama, Sheikh Iqbal Ahamed

**Affiliations:** Ubicomp Lab, Department of Computer Science, Marquette University, Wisconsin, USA

## Abstract

Pancreatic ductal adenocarcinoma (PDAC) is one of the most lethal cancers worldwide. It has a five-year survival rate of approximately 13%, largely due to late diagnosis, rapid metastasis, and limited therapeutic options. The tumor microenvironment (TME), including immune and non-immune cells, plays a role in PDAC progression and drug resistance. However, molecular characteristics of PDAC based on TME-based subtypes are not well studied. To address this, we developed a multi-layered Graph Convolutional Network (GCN) to identify TME-guided PDAC subtypes. First, we constructed a TME reference atlas using single-cell RNA-seq data comprising immune and non-immune cell types. Second, we applied in silico cytometry to estimate cell-type proportions in bulk tumor RNA-seq samples from the Cancer Genome Atlas PDAC patients. Third, we constructed the GCN model integrating the estimated patient-specific cellular proportions with multi-omics datasets, such as gene expression, DNA methylation, copy number variation, and clinical data. Finally, we predicted PDAC subtypes using an unsupervised clustering method on the GCN-based embedding matrices. Our method identified seven PDAC subtypes, which we further grouped into two ecological tumor families. These subtypes exhibited distinct immune infiltration, stromal activity, and malignant epithelial cell abundance. Differential gene expression analysis revealed subtype-specific molecular programs. Survival analysis demonstrated significant differences in overall survival across ecological families and subtypes, highlighting their prognostic relevance. Thus, our method provides a biologically informed framework for patient stratification, with potential implications for advancing precision oncology approaches.

## Background

PDAC is an aggressive cancer characterized by molecular, cellular, and microenvironmental heterogeneity. Using bulk sequencing transcriptome, molecular subtypes of PDAC were identified^1^. However, a defining feature of this cancer is its dense and heterogeneous tumor microenvironment (TME). It contains diverse populations of cancer-associated fibroblasts (CAFs), myeloid cells, lymphocytes, endothelial cells, and other stromal populations^2,3^. The PDAC TME was found to influence tumor growth, metabolic adaptation, and immune evasion^4,5^. The desmoplastic stroma and immunosuppressive TME characteristic of PDAC are also considered important contributors to its resistance to chemotherapy and immune-based therapies^5,6^.

Single-cell RNA sequencing (scRNA-seq) has substantially refined our understanding of the TME by characterizing transcriptionally distinct malignant, immune, and stromal populations that are obscured in bulk tumor profiles. Early single-cell studies of human PDAC showed inter- and intra-tumoral heterogeneity in both malignant and non-malignant cells^7^. A recent study generated scRNA-seq datasets, GSE194247 and GSE235449, that provide profiles of non-immune (17 samples) and immune (3 samples) cell populations, respectively, from pancreatic tumors^8^. This study identified five distinct functional subclusters of pancreatic cancer cells and six distinct cancer-associated fibroblast subclusters. Despite the high cellular resolution provided by scRNA-seq, single-cell cohorts generally contain substantially fewer samples than large bulk transcriptomic cohorts. Furthermore, scRNA-seq studies are limited by a lack of matched multi-omics and clinical information. On the other hand, bulk sequencing projects, such as The Cancer Genome Atlas (TCGA)^9^, provide genomic, transcriptomic, epigenomic, and clinical characterization across a large patient cohort (~200 PAAD samples).

Computational deconvolution provides a strategy for using single-cell profiles to estimate the cellular composition of heterogeneous bulk tissues. *MuSiC*, for example, uses scRNA-seq reference information to estimate cell-type proportions from bulk RNA-seq while accounting for cross-subject variation^10^. *CIBERSORTx* extended digital cytometry by enabling single-cell-derived reference signatures to be applied to bulk transcriptomes and allowing inference of both cell abundance and cell-type-specific expression profiles^11^. More recently, *BayesPrism* introduced a Bayesian framework for using scRNA-seq references to deconvolute bulk tumor transcriptomes while accounting for differences between reference and bulk expression profiles^12^. These approaches establish a conceptual and methodological bridge through which cellular information derived from small single-cell cohorts can be projected onto larger collections of bulk tumors.

Conventional unsupervised clustering approaches, including hierarchical and consensus clustering, have been instrumental in identifying molecular cancer subtypes, particularly in establishing transcriptomic classifications of PDAC^1,13,14^. Nevertheless, feature-based clustering does not explicitly represent relationships among individual patients. Patient similarity networks provide an alternative representation in which patients are modeled as nodes and edges encode similarity in their molecular or phenotypic profiles. This network-based representation is particularly suitable for integrating multi-omics data because different omics layers capture distinct dimensions of tumor biology. Copy-number variation (CNV) reflect genomic changes, DNA methylation represents an important layer of epigenomic regulation, and mRNA captures transcriptional profiles. Importantly, patients that are highly similar according to one modality may not necessarily be the most similar according to another. Graph neural networks enable machine-learning models to learn directly from network data. Graph convolutional networks (GCNs) generate node embeddings that allow the learned representation of an individual node to incorporate both its own characteristics and information from its local graph context^15^. Graph learning has increasingly been applied to multi-omics cancer analysis. MOGONET constructed omics-specific sample-similarity graphs and used GCNs to integrate information across multiple omics modalities for biomedical classification^16^. MoGCN subsequently combined dimensionality-reduced multi-omics features with patient similarity networks and GCN learning for cancer subtype analysis^16^. Nevertheless, most existing graph-based cancer stratification frameworks have primarily constructed patient relationships from bulk tumor-centric multi-omics data. Comparatively less attention has been given to single-cell-informed TME states as a primary source of patient similarity. Single-cell-informed deconvolution provides an opportunity to transform the stromal and immune contributions within bulk expression data into patient-level features. A TME-derived patient graph could therefore organize patients according to similarities in their cellular ecosystems, while genomic, epigenomic, transcriptomic, and clinical graphs could represent complementary dimensions of tumor-intrinsic molecular heterogeneity.

In this study, we developed a multi-omics-based GCN framework to predict TME-driven subtypes in PDAC. We first constructed a PDAC-specific cellular reference representing complementary immune and non-immune cell populations from GSE194247 and GSE235449 datasets (**Fig. 1; Methods**). Cell-type and cell-state expression signatures derived from this atlas were then used to deconvolute PDAC bulk RNA-seq data from TCGA and generate quantitative patient-level TME profiles. These profiles were subsequently considered together with gene expression, CNV, DNA methylation, and clinical features of the TCGA PDAC samples. We then constructed a multi-view patient graph in which patients were modeled as nodes and edges captured similarities in patients’ characteristics. Using a GCN architecture, we sought to generate low-dimensional patient embeddings that preserve both patient-specific features and higher-order relationships among tumors. Finally, we applied unsupervised clustering to these graph-derived representations to predict TME-driven and multi-omics-refined patient subtypes (**Fig. 1; Methods**).

**Figure 1:**
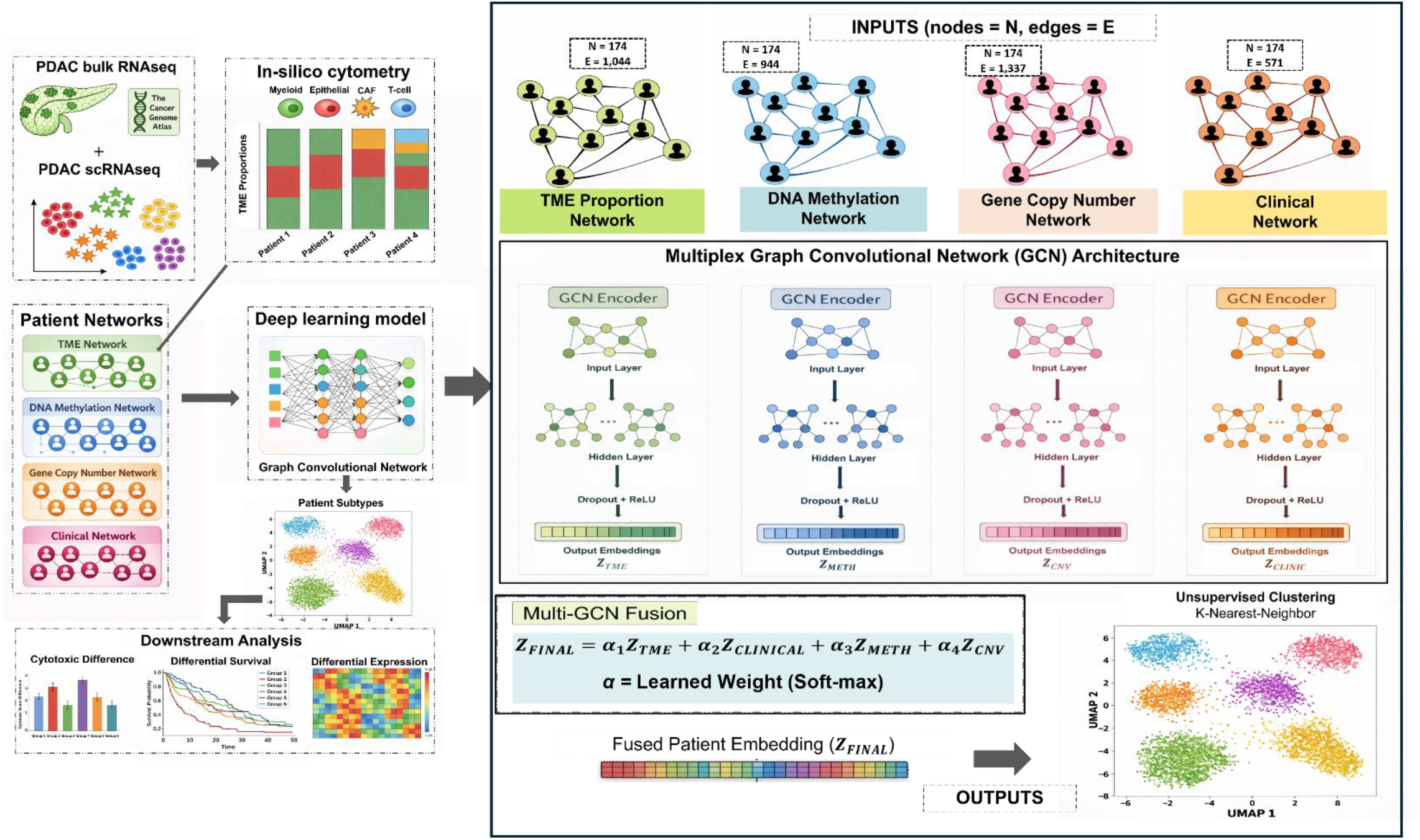
Overview of the proposed multi-omics GCN framework for PDAC patient stratification.

## Results

### We predicted seven distinct TME-associated PDAC subtypes

Our multiplex-GCN framework generated integrated patient embeddings, which were subjected to unsupervised clustering that identified seven distinct PDAC patient subtypes (**Methods; Fig. 2**). The identified subtypes achieved an overall silhouette score of 0.41 in the learned GCN embedding space, indicating meaningful separation among patient groups. For visualization, the patient embeddings were projected into two dimensions using UMAP, with patients colored according to their assigned subtypes (**Fig. 2**). We next investigated whether these GCN-derived subtypes were associated with distinct TME compositions. For this, we compared the centered log-ratio (CLR)-transformed mean cell-type abundances across the seven PDAC subtypes. We observed substantial variation in the proportions of myeloid cells, CAFs, malignant epithelial cells, plasma cells, T cells, and B cells (**Fig. 3**). These differences indicate that the identified patient groups are characterized by distinct cellular compositions within the tumor microenvironment.

**Figure 2:**
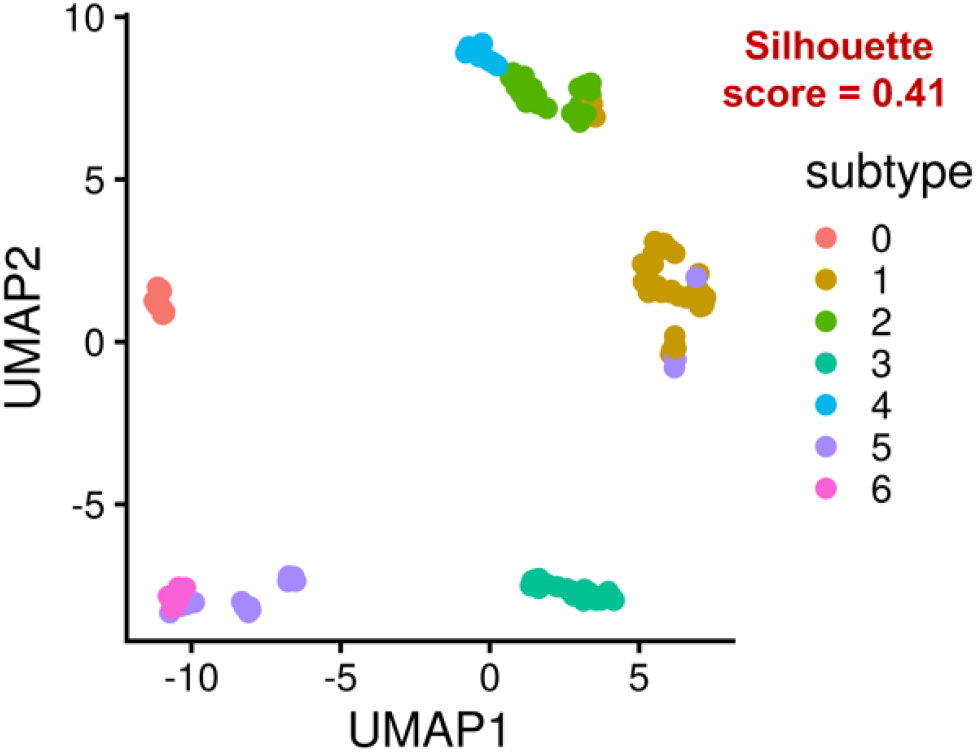
Two-dimensional UMAP visualization of patient embeddings learned by the TME-guided multiplex GCN. Each point represents one patient and is colored according to the assigned subtype (0-6).

**Figure 3:**
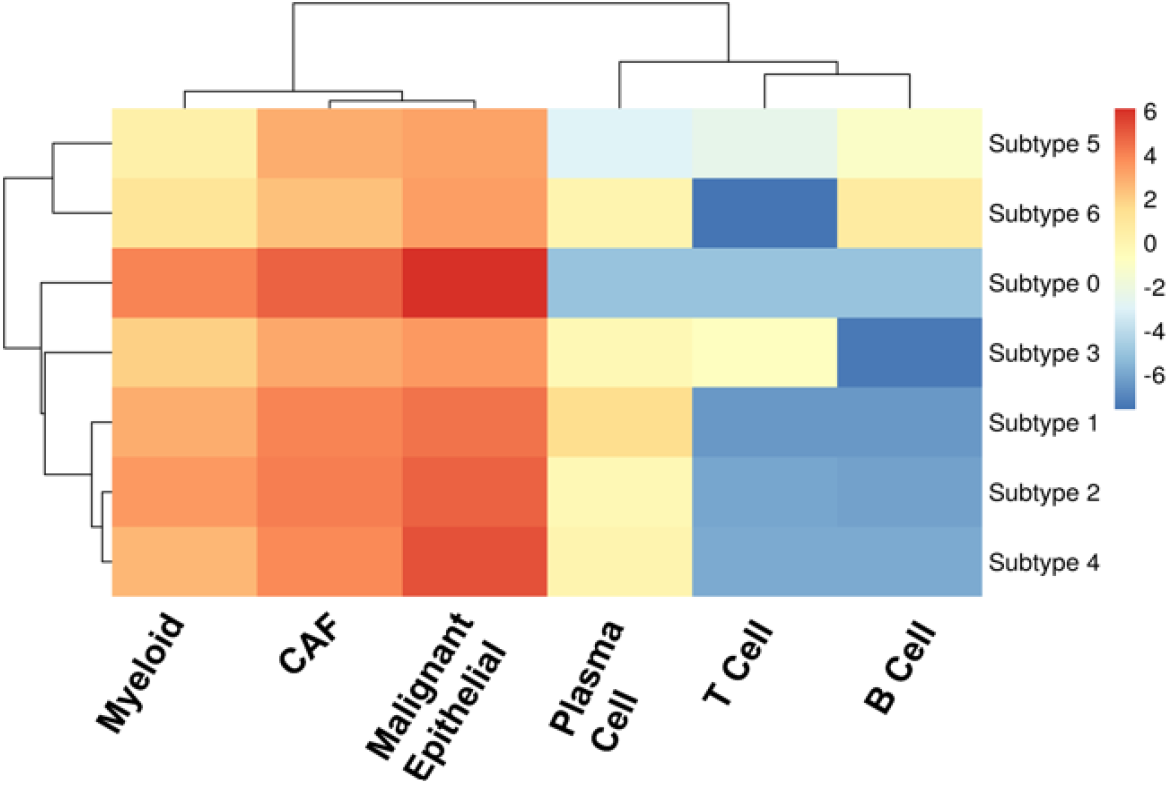
Heatmap showing subtype-level CLR-transformed abundances of six TME cell types. Hierarchical clustering illustrates differences in TME composition across the seven PDAC subtypes and the higher-order organization of the identified patient groups.

To further characterize the biological differences among the identified PDAC subtypes, we curated a panel of genes representing epithelial differentiation, mesenchymal and stromal signaling, inflammatory responses, cytotoxic immune activity, and tumor-associated pathways (**Supplementary File X**). Using PDAC bulk RNA-seq data, we computed the median expression of each gene across patient samples within each subtype and generated a hierarchically clustered heatmap of gene-wise standardized expression profiles (**Fig. 4**). We selected *EPCAM* and *KRT19* to represent the epithelial program because they are well-established markers of epithelial cell identity and differentiation^7^. The cytotoxic program was defined using *GZMB* and *IFNG*, which are associated with cytotoxic lymphocyte effector functions and immune activation^17,18^, together with *CXCL9* and *CXCL10*, which are interferon-inducible chemokines involved in immune-cell recruitment^19^. The analysis revealed distinct transcriptional patterns across the seven subtypes, including differences in epithelial markers (*EPCAM* and *KRT19*) and immune-related genes (*IFNG, GZMB, CXCL9*, and *CXCL10*). The transcriptional profiles revealed two broader ecological families previously identified from TME cellular composition, comprising subtypes 0-4 and subtypes 5-6. However, individual subtypes also exhibited distinct transcriptional characteristics. For example, subtype 4 showed relatively elevated expression of epithelial markers, whereas subtype 3 exhibited increased expression of cytotoxic and immune-activation genes. Together, these findings suggest that our method captures transcriptional heterogeneity at both the subtype and broader ecological levels.

**Figure 4:**
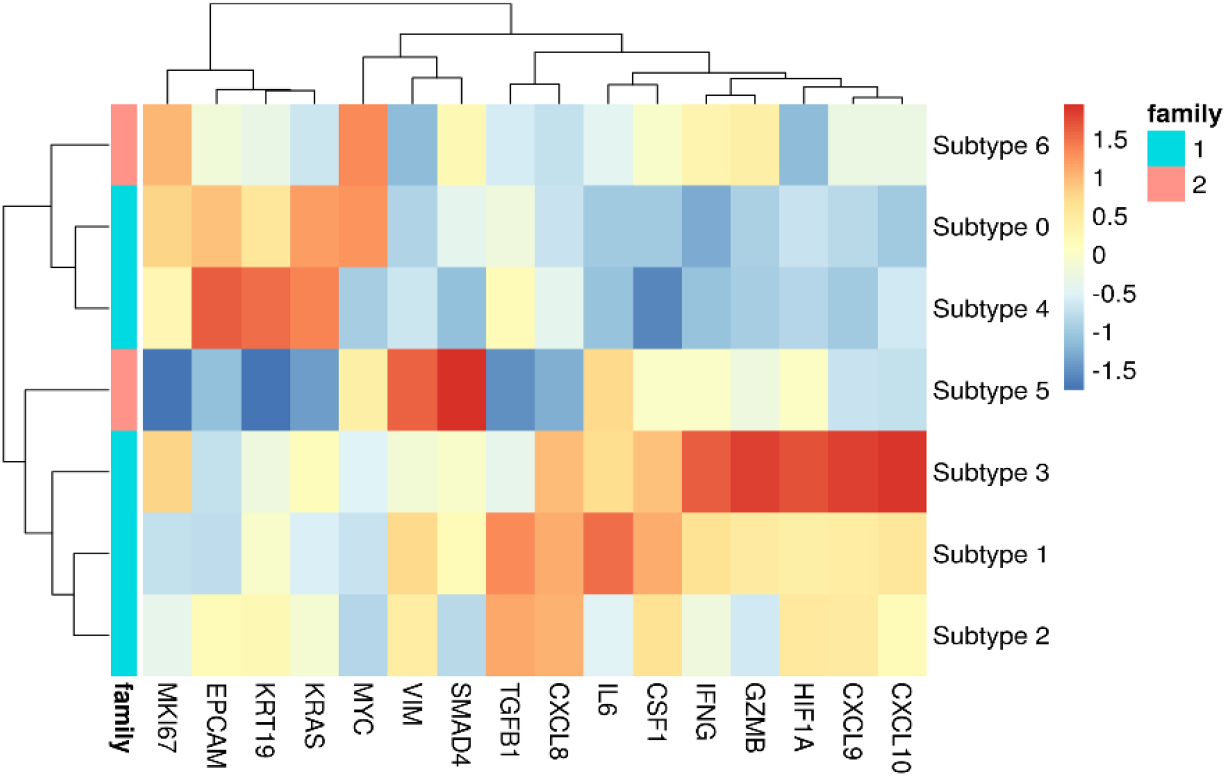
Heatmap showing subtype-level expression patterns of representative genes associated with tumor-intrinsic, epithelial, stromal/TGF-β, inflammatory/myeloid, cytotoxic/immune, hypoxia, and proliferation programs. Expression values were summarized at the subtype level and standardized by gene (z-score) for visualization. Hierarchical clustering demonstrates distinct transcriptional states across the seven PDAC subtypes. The annotation indicates membership in the two higher-order ecological families.

To further quantify transcriptional differences among the identified PDAC subtypes, we calculated patient-level epithelial and cytotoxic program scores using PDAC bulk RNA-seq data (**Fig. 5**). For each patient, program scores were calculated as the mean expression of the corresponding genes. We compared these scores between the two ecological families using Wilcoxon rank-sum tests and across the seven individual subtypes using Kruskal–Wallis tests.

**Figure 5.**
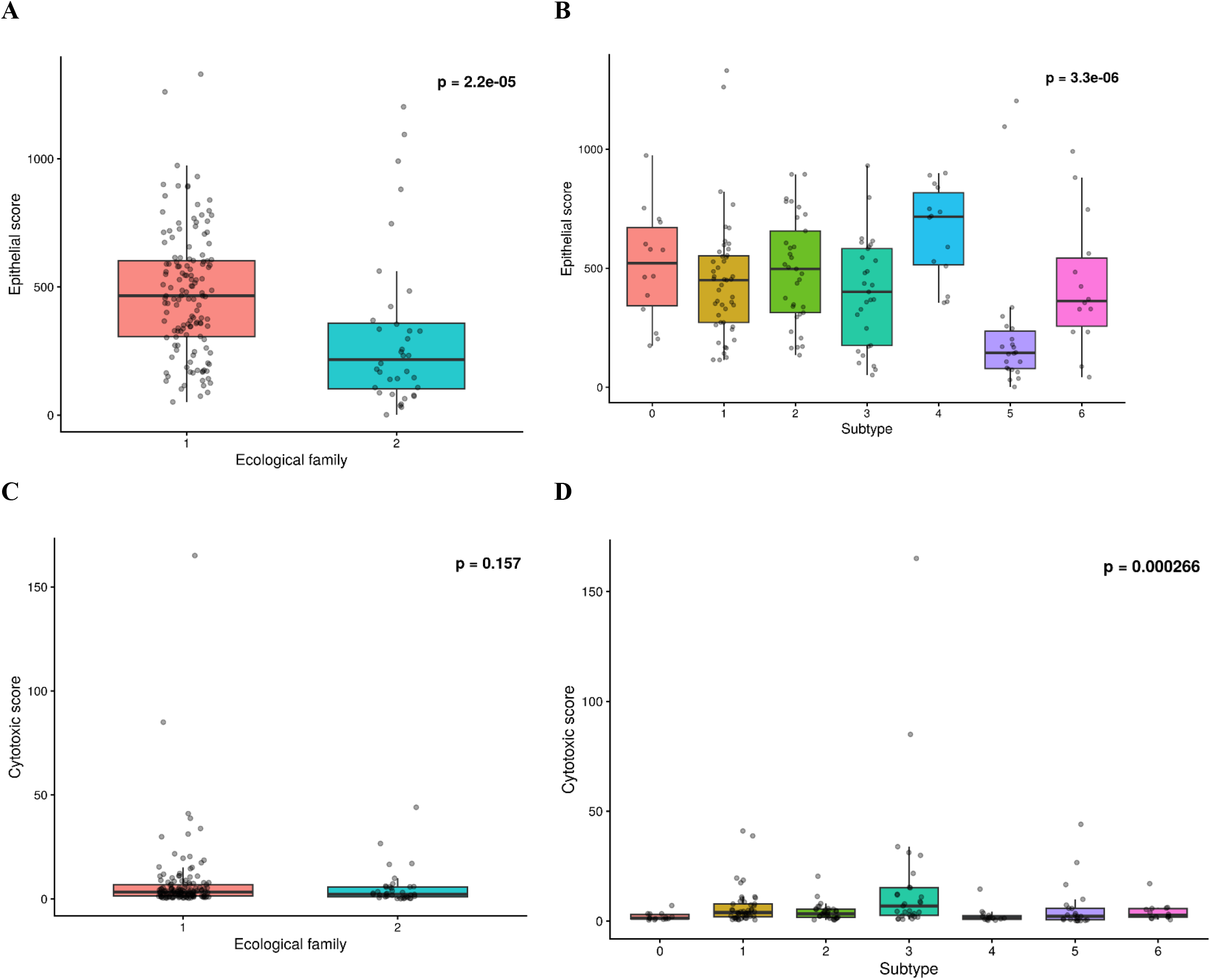
Epithelial and cytotoxic transcriptional programs differ across PDAC ecological families and subtypes. **(A)** Epithelial program scores in ecological Family 1 and Family 2 (Wilcoxon rank-sum *P* = 2.2 × 10. ^−5^); **(B)** Epithelial program scores across the seven graph-derived PDAC subtypes (Kruskal–Wallis *P* = 3.3 × 10^−6^); **(C)** Cytotoxic program scores between ecological families (*P* = 0.157; **(D)** Cytotoxic program scores across the seven subtypes (Kruskal–Wallis *P* = 2.66 × 10^−4^).

At the ecological-family level, epithelial program scores were significantly higher in the family comprising subtypes 0-4 than in the family comprising subtypes 5-6 (Wilcoxon *P* = 2.2 × 10^−5^; **Fig. 5A**). In contrast, cytotoxic program scores did not differ significantly between the two families (*P* = 0.157; **Fig. 5B**). At the individual-subtype level, however, both transcriptional programs exhibited significant heterogeneity. Epithelial program scores differed significantly across the seven subtypes (Kruskal–Wallis *P* = 3.3 × 10^−6^; **Fig. 5C**), with subtype 4 exhibiting the highest score and subtype 5 the lowest. Similarly, cytotoxic program scores varied significantly across subtypes (Kruskal–Wallis *P* = 2.66 × 10^−4^; **Fig. 5D**), with subtype 3 showing the strongest cytotoxic transcriptional profile. Notably, although cytotoxic program scores did not differ significantly between the two broader ecological families, their significant variation across individual subtypes suggests that subtype-specific immune transcriptional differences may be obscured when patients are grouped into broader families. Collectively, these findings support a hierarchical organization of PDAC heterogeneity, in which broader TME-associated ecological families coexist with finer subtype-specific epithelial and immune transcriptional states.

### PDAC subtypes and ecological families exhibit overall survival differences

We next examined whether the predicted subtypes were associated with differences in clinical outcomes in terms of overall survival of patients. Kaplan-Meier survival curves were generated using the R package *survival*^20^, and differences between groups were assessed using log-rank tests. Patients in ecological family 2 showed substantially better overall survival than those in ecological family 1 (log-rank *P* = 5.7 × 10^−4^; **Fig. 6A**). Overall survival also differed significantly across the seven individual PDAC subtypes (log-rank *P* = 0.0043; **Fig. 6B**). Together, these findings demonstrate the clinical relevance of the identified PDAC patient stratification.

**Figure 6:**
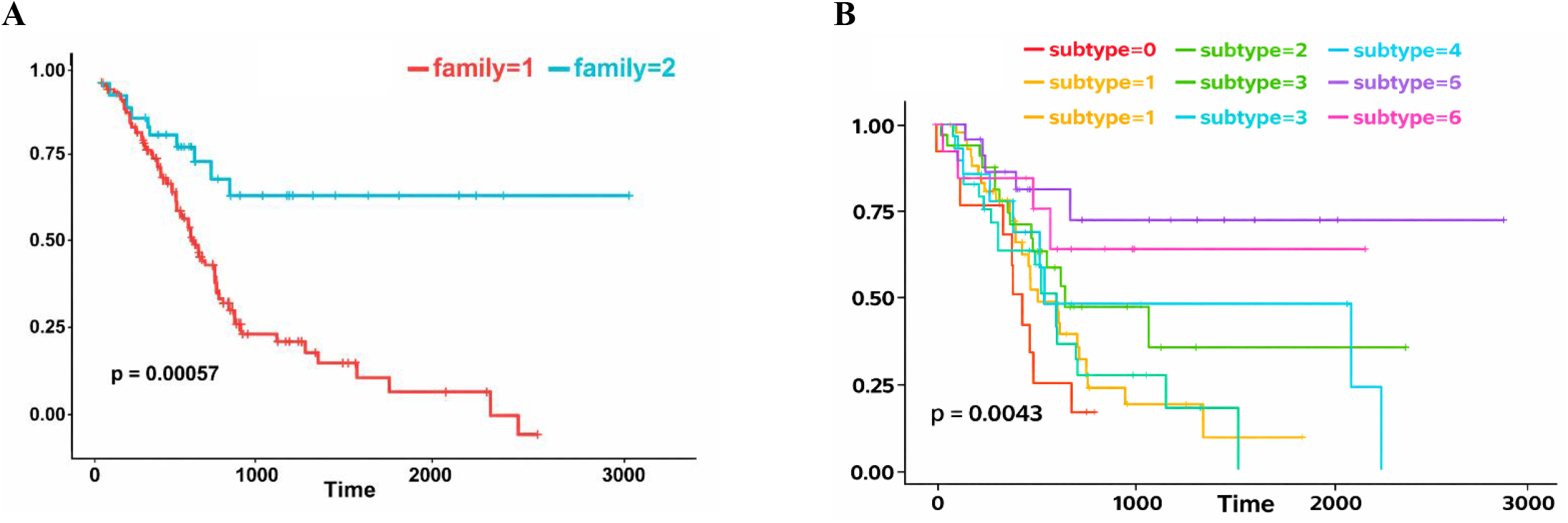
Kaplan-Meier analysis of overall survival according to **(A)** the two ecological families (log-rank *P* = 5.7 × 10^−4^) and **(B)** the seven graph-derived PDAC subtypes (log-rank *P* = 0.0043).

**Figure 7:**
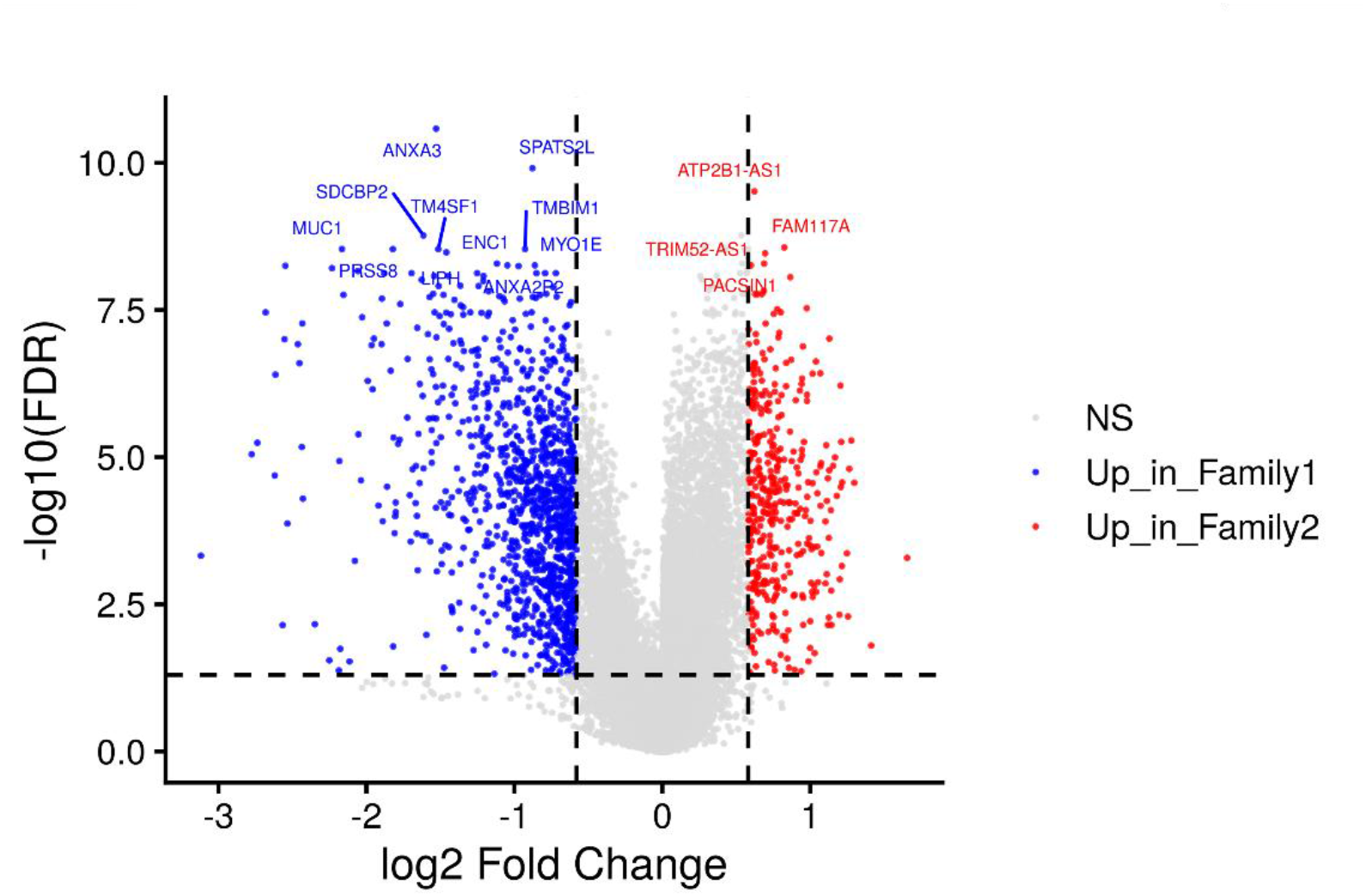
Volcano plot showing genome-wide differential expression between ecological Family 2 and Family 1. The x-axis represents log2 fold change (Family 2 versus Family 1), and the y-axis represents −log10(FDR-adjusted *P* value). Differentially expressed genes were identified using FDR < 0.05; an additional threshold of |log2FC| ≥ 0.58 was used to highlight genes with appreciable expression differences. Selected highly significant genes are labeled.

### Ecological families exhibit differential gene expression patterns

To characterize the molecular differences in the predicted patient groups, we performed genome-wide differential gene expression analysis comparing ecological Family 2 (subtypes 5-6) with Family 1 (subtypes 0-4) using PDAC bulk RNA-seq data (**Methods**). We used the R package *limma*^21^, with expression values averaged across aliquots at the patient level and log2(x + 1)-transformed. Gene-wise linear models were fitted with ecological family as the explanatory variable, followed by empirical Bayes variance moderation. Differentially expressed (DE) genes were identified using a Benjamini-Hochberg-adjusted false discovery rate (FDR) < 0.05 and an absolute log2 fold-change threshold of ≥ 0.58. Applying these criteria, we identified 1,535 DE genes, including 1,151 genes with higher expression in Family 1 and 384 genes with higher expression in Family 2 (**Fig. 6**). Positive log2 fold changes indicated higher expression in Family 2, whereas negative values indicated higher expression in Family 1.

Among the most significant differentially expressed genes, *ANXA3, SDCBP2, MUC1, PRSS8, TM4SF1, ENC1, SPATS2L, TMBIM1*, and *MYO1E* showed higher expression in Family 1, whereas *ATP2B1-AS1, TRIM52-AS1, PACSIN1*, and *FAM117A* showed higher expression in Family 2 (**Fig. 6**). Together, these findings demonstrate extensive transcriptional differences between the two TME-associated ecological families.

### Pathway enrichment reveals functional differences between ecological familie

To investigate the biological processes underlying transcriptional differences between the two ecological families, we performed pathway enrichment analyses using the R package *clusterProfiler*^22^. We obtained gene annotations from *org*.*Hs*.*eg*.*db*^23^ and Hallmark gene sets retrieved from MSigDB^24^ using *msigdbr*^25^. Gene Set Enrichment Analysis was performed using the complete gene list ranked by log2 fold changes between Family 2 and Family 1. The analysis identified 35 significantly enriched Hallmark pathways (adjusted *P* < 0.05; **Fig. 8A**). Among these, pancreas beta cell and spermatogenesis gene signatures were positively enriched in Family 2, whereas epithelial–mesenchymal transition, TNFα signaling via NF-κB, and interferon-α response were enriched in Family.

**Figure 8:**
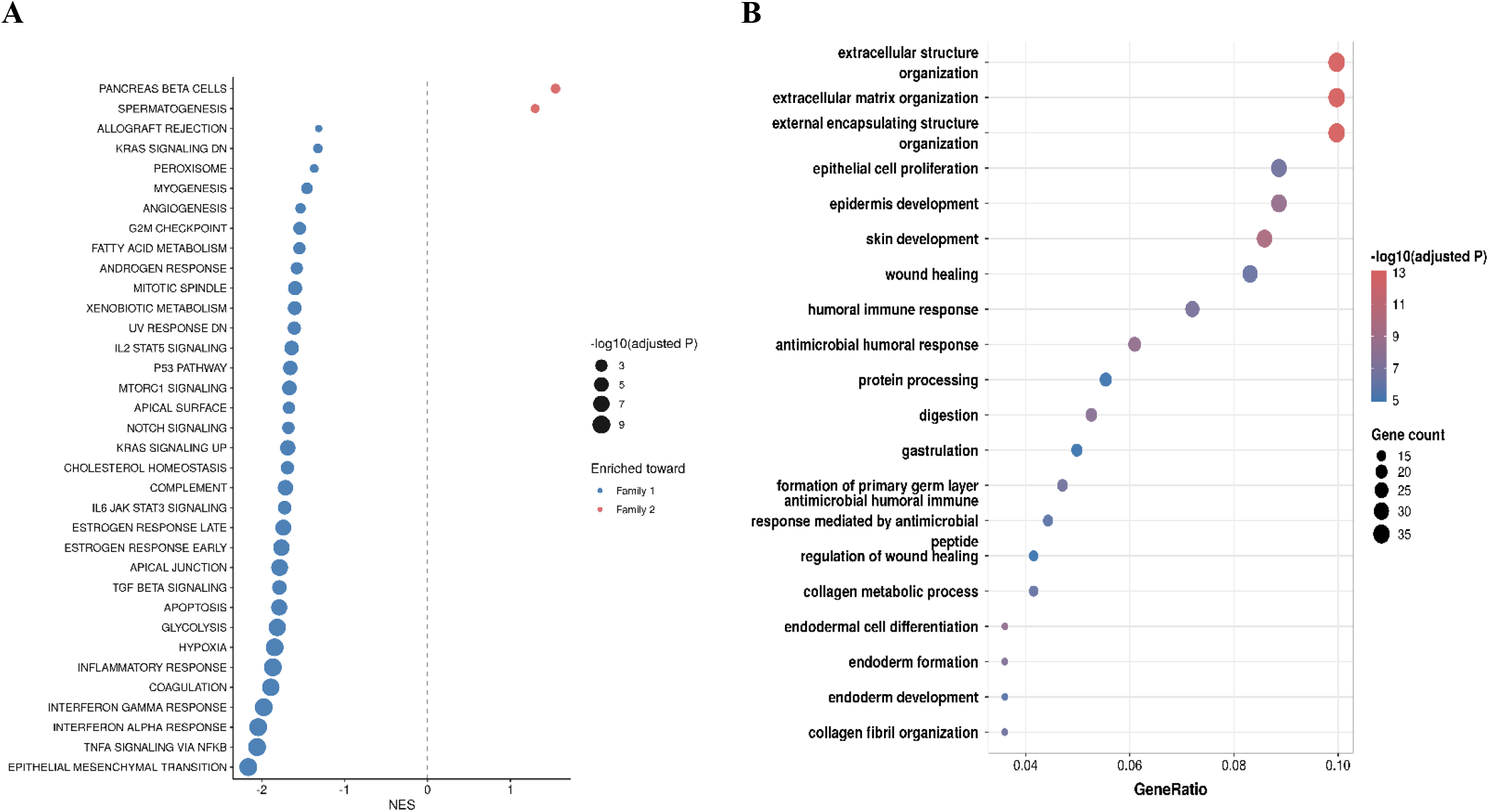
Dot plots showing (A) Hallmark GSEA pathways enriched in Family 1 (negative NES, blue) and Family 2 (positive NES, red), and (B) the top 20 enriched GO Biological Process terms based on differentially expressed genes. Dot size represents statistical significance in (A) and gene count in (B).

To further characterize functional differences, we performed Gene Ontology (GO)^26^ Biological Process enrichment analysis using the over-representation analysis approach implemented in *clusterProfiler*, selecting DE genes with FDR < 0.05 and |log2FC| > 1. Enrichment significance was assessed using Benjamini-Hochberg-adjusted P values. This analysis identified significantly enriched biological processes, including extracellular matrix organization, extracellular structure organization, and wound healing (**Fig. 8B**). These findings suggest that the two predicted ecological families exhibit distinct biological programs involving extracellular matrix remodeling, inflammatory and immune responses, and epithelial-mesenchymal transition, providing functional context for the transcriptional heterogeneity identified through genome-wide differential expression analysis.

### Model sensitivity analyses support the contribution to predicted subtypes

To assess the robustness of the identified PDAC subtypes, we evaluated whether our prediction remained consistent under alternative model settings. These analyses examined the effects of changes in model initialization, training parameters, and the relative contributions of different molecular and clinical data modalities. Across most configurations, the model assigned the greatest fusion weight to TME, whereas clinical characteristics, DNA methylation, and CNV received substantially smaller weights (**Supplementary Fig. S1**). The selected model achieved the highest silhouette score of approximately 0.41, while several alternative configurations maintained the scores between 0.35 and 0.40 (**Supplementary Fig. S2**). This indicates relatively consistent separation of patient groups in different settings. In contrast, configurations that reduced the influence of TME information or changed how the model combined the different data types showed weaker separation of patient groups, with silhouette scores as low as approximately 0.19. Together, these findings suggest that the identified patient stratification is relatively robust to several modeling choices, while highlighting the important contribution of TME in the subtype prediction.

## Methods

We developed a TME-guided multiplex network-based GCN framework to identify patient subtypes in PDAC samples from the TCGA database.

### Network Creation

We constructed four different patient-similarity networks representing TME composition, clinical characteristics, DNA methylation, and CNV using custom Python scripts with *NumPy*^27^, *pandas*^28^, *PyTorch*^29^, and *scikit-learn*^30^. In each network, nodes represented patients, and weighted edges represented modality-specific similarities.

The TME network was constructed using estimated fractions of T cells, B cells, plasma cells, myeloid cells, CAFs, and malignant epithelial cells. To account for the compositional nature of these data, cell-type fractions were transformed using the CLR transformation with a pseudocount of e10-5, and patient similarities were calculated from Aitchison distances using a radial basis function (RBF) kernel.

The clinical network incorporated eight variables: age at diagnosis, sex, race, smoking history, diabetes history, chronic pancreatitis history, alcohol history, and tissue source site. Missing categorical values were assigned an “Unknown” category, smoking history was classified as never or ever smoking (with unknown status retained), and age was Z-score standardized. Categorical variables were one-hot encoded to generate patient-level features for the GCN, while 1-Gower distance was used to calculate pairwise patient similarities.

For the DNA methylation and CNV networks, we used gene-level methylation beta values and copy-number variation profiles, respectively. For each modality, the 1,000 most variable genes were selected based on median absolute deviation (MAD), followed by Z-score standardization. Pairwise patient similarities were then calculated using Pearson correlation coefficients. Additional graph sparsification and degree-based neighborhood pruning were applied as specified in **Table 1**. Patients were harmonized across all four modalities, resulting in a final cohort of 174 patients with matched network nodes. Isolated nodes were retained to maintain cohort consistency. The network construction parameters, feature dimensions, and final network characteristics are summarized in **Table 1**.

**Table 1.** Network construction parameters and final characteristics of the four patient-similarity layers of 174 patients. Degree-based neighborhood pruning values represent the number of highest-weight connections selected per patient. Isolated nodes were retained to maintain consistent patient representation across the four network layers. NA means not applied.

| Parameter | Clinical | TME | DNA methylation | CNV |
| --- | --- | --- | --- | --- |
| Final node features | 41 | 6 | 1,000 | 1,000 |
| kNN (k) | 5 | 10 | 12 | 12 |
| Target average degree for global edge sparsification | NA | 12 | 14 | NA |
| Degree-based neighborhood pruning | NA | NA | 25 | 30 |
| Final number of edges | 571 | 1,044 | 944 | 1,337 |
| Isolated nodes (n) | 0 | 4 | 19 | 0 |

### Network-specific graph convolutional encoding

We developed an unsupervised TME-guided multiplex GCN architecture to learn integrated patient representations from four patient-similarity networks: TME composition, clinical characteristics, DNA methylation, and CNV. Our model comprised three main components: (i) network-specific graph convolutional encoding, (ii) learnable TME-guided embedding fusion, and (iii) unsupervised graph reconstruction for representation learning, followed by patient subtype discovery using K-means clustering (**Fig. 1**).

We represented each patient-similarity network as a weighted graph:

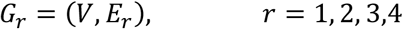

where *V* represents the common set of 174 patients, and *E*_*r*_ denotes the weighted patient–patient connections in the network *r*. We represented each network using a weighted adjacency matrix *A*_*r*_.

We added self-loops to each adjacency matrix to preserve patient-specific information during graph convolution:

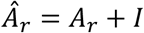

We then computed the symmetrically normalized adjacency matrix:

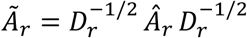

where *I* is the identity matrix, *D*_*r*_ is the diagonal degree matrix calculated from *Â*_*r*_, and 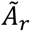 is the normalized adjacency matrix.

We constructed four separate two-layer GCN encoders, each operating on a distinct patient-similarity network. In the first graph convolutional layer, we transformed the input patient features and aggregated information from neighboring patients. We subsequently applied a rectified linear unit (ReLU) activation function and dropout:

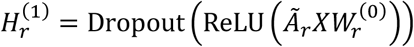

We then applied a second graph convolutional layer to generate the network-specific patient embeddings:

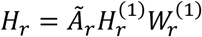

Here, *X* denotes the input patient-feature matrix, 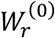 and 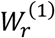 represent network-specific trainable weight matrices, 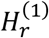 denotes the hidden representation, and *H*_r_ represents the final network-specific patient embeddings.

We used a hidden dimension of 64 and an output embedding dimension of 32. Consequently, each GCN encoder generated a 32-dimensional patient representation that captured information propagated through its corresponding network structure.

Next, we defined an integrated patient embedding matrix to combine the representations generated by the four GCN encoders as:

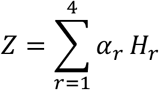

where *Z* represents the integrated patient embedding matrix, *H*_*r*_ denotes the embeddings generated from network *r*, and *α*_*r*_ represents the learned contribution of that network.

We introduced trainable fusion parameters and normalized them using a softmax function:

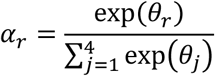

Where *θ*_*r*_ denotes the trainable fusion parameter associated with network *r*. This normalization ensured that the fusion weights were nonnegative and summed to one.

To guide the integrated representation toward TME-associated heterogeneity, we introduced a predefined prior distribution for the four network contributions:

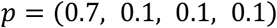

These values corresponded to the TME, clinical, DNA methylation, and CNV networks, respectively.

Instead of fixing the fusion weights, we regularized their learned values toward the prior distribution using Kullback– Leibler (KL) divergence:

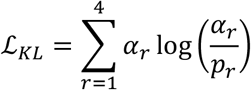

where *p*_*r*_ represents the predefined prior weight for network *r*. This regularization encouraged greater contribution from the TME network while allowing the model to learn the relative importance of clinical and molecular information.

### Training predictive models using node embedding combinations

We trained the multiplex GCN without predefined patient subtype labels using a graph link-reconstruction objective. For each patient-similarity network, we considered observed patient–patient connections as positive examples and randomly sampled an equal number of unconnected patient pairs as negative examples.

We computed the probability of a connection between patients *i* and *j* using a dot-product decoder applied to their integrated embeddings:

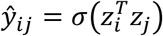

Where *z*_*i*_ and *z*_*j*_ represent the integrated embedding vectors of patients *i* and *j*, respectively, and *σ* denotes the sigmoid activation function.

We calculated the graph reconstruction loss for each network using binary cross-entropy:

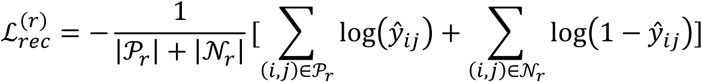

Where *P*_*r*_ represents the observed patient connections and *N*_*r*_ represents the sampled non-connections for network *r*.

By reconstructing patient relationships across the four network layers using the shared integrated embeddings, we encouraged the model to preserve complementary patient-similarity patterns across modalities.

We constructed a joint optimization objective that combined the reconstruction losses from all four networks with the TME-guided KL divergence regularization:

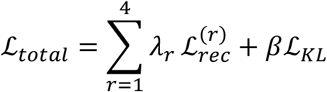

where *λ*_*r*_ represents the reconstruction-loss weight for network *r*, and *β* controls the contribution of the KL regularization term.

To prioritize TME-associated patient relationships, we assigned a reconstruction-loss weight of 8 to the TME network and a weight of 1 to each of the clinical, DNA methylation, and CNV networks. Therefore, our final optimization objective was:

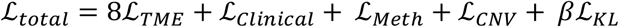

We set the KL regularization coefficient to 1.0 and optimized the model using the Adam optimizer with a learning rate of 0.01 and weight decay of 10^−4^. We used a hidden dimension of 64, a final embedding dimension of 32, and dropout of 0.3. We trained the model for 300 epochs.

Following model optimization, we extracted the 32-dimensional integrated patient embeddings for all 174 patients. These embeddings represented the learned patient relationships across the multiplex network and were subsequently used for unsupervised patient subtype discovery.

Following GCN training, we extracted the integrated 32-dimensional patient embeddings to identify patient subtypes without using predefined subtype labels. We applied K-means clustering to the learned embeddings to group patients with similar integrated network representations.

We evaluated candidate cluster numbers ranging from 2 to 8 and computed the silhouette score for each clustering solution. The silhouette score measures how closely patients are grouped within their assigned cluster relative to patients in other clusters. For each patient *i*, we calculated the silhouette coefficient as:

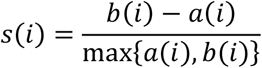

where *a*(*i*) represents the average distance between patient *i* and other patients within the same cluster, and *b*(*i*) represents the minimum average distance between patient *j* and patients in any other cluster.

We then computed the overall silhouette score by averaging the individual silhouette coefficients across all patients:

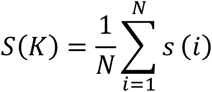

Where *N* represents the total number of patients and *K* denotes the number of clusters.

We selected the optimal number of patient subtypes by identifying the clustering solution with the highest average silhouette score:

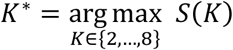

where *K*^∗^ represents the optimal number of clusters.

Finally, we assigned each patient to the corresponding cluster based on the selected K-means solution. We considered these clusters as the newly identified patient subtypes and used the resulting subtype assignments for downstream biological and clinical characterization.

Following model training, the integrated patient embeddings were subjected to unsupervised K-means clustering. Candidate solutions containing two through eight clusters were evaluated. K-means was run with 50 initializations for each candidate number of clusters, and the optimal number of clusters was selected according to the highest silhouette score calculated in the learned embedding space.

This procedure identified seven PDAC subtypes (subtypes 0-6). For visualization, the learned patient embeddings were projected into two dimensions using UMAP with 15 nearest neighbors, a minimum distance of 0.10, and cosine distance. The reported subtype silhouette score was calculated from the original learned embedding rather than the two-dimensional UMAP representation.

### Building PDAC reference panel from scRNA-seq datasets to estimation of patient-level TME composition

We downloaded raw 10x Genomics scRNA-seq count matrices for immune (**GSE194247**) and non-immune cells (**GSE235449**) from the GEO database^8^. We used the *Seurat*^31^ R package to filter these matrices, requiring genes to be detected in at least three cells and cells to contain at least 200 detected features. For the non-immune dataset, cells with >400 and <6,500 detected genes and <20% mitochondrial transcripts were retained. For the immune dataset, cells with >300 and <6,000 detected genes and <20% mitochondrial transcripts were retained. The distributions of detected features, total RNA counts, and mitochondrial transcript percentages before and after filtering were examined to assess the effects of quality-control filtering (**Supplementary Fig. S3-S6**). Seurat *SCTransform* was applied separately to each dataset, 3,000 integration features were selected, integration anchors were identified using *SCT* normalization, and the datasets were integrated using *IntegrateData*. Principal-component analysis was performed using 30 principal components, followed by UMAP visualization, nearest-neighbor graph construction, and clustering at a resolution of 0.4 to assess integration and cellular structure.

We utilized curated marker-gene sets [] representing immune and non-immune PDAC cell populations to assign cell identities. We removed the marker sets containing fewer than three genes. Cell-type-specific marker scores were calculated using Seurat’s *AddModuleScore* function, and each cell was assigned to the cell type corresponding to its highest module score. Annotation confidence was evaluated by calculating the difference between the highest and second-highest module scores, and cells with a score difference below 0.2 were excluded from subsequent reference construction.

Cell types were consolidated into six TME populations: T cells, B cells, plasma cells, myeloid cells, CAFs, and malignant epithelial cells. A pseudobulk transcriptional reference was constructed from the annotated single-cell data to reduce the influence of single-cell sparsity and stochastic dropout. Raw RNA counts were summed across cells assigned to each of the six final cell populations. The resulting cell-type expression profiles were normalized to counts per million (CPM), producing a six-component PDAC-specific pseudobulk transcriptional reference. The correlation structure was examined to evaluate transcriptional relationships among the reference populations before their use in bulk-tumor deconvolution (**Supplementary Fig. S7**). We utilized the single-cell reference and PDAC bulk transcriptomic profiles with *CIBERSORTx*^32^ to estimate the patient-level relative abundance of the six TME cell types. Because estimated cell fractions represent compositional data, CLR transformation was applied before TME-based similarity analysis.

## Discussion

In this study, we developed a TME-guided GCN framework to characterize heterogeneity among patients with PDAC. By integrating patient-similarity networks representing TME composition, clinical characteristics, DNA methylation, and CNV, the framework predicted seven PDAC subtypes with distinct microenvironmental and transcriptional characteristics. These subtypes were further grouped into two consensus ecological families that differed in gene-expression patterns and overall survival. Model sensitivity analyses showed that patient subtype prediction remained relatively stable across multiple model configurations, while reducing the contribution of the TME substantially weakened subtype separation. Together, these findings support the concept that the cellular organization of the PDAC microenvironment represents an important dimension of interpatient heterogeneity that can complement conventional molecular and clinical features.

We computed a PDAC-specific single-cell atlas with immune and non-immune cell types and applied it to bulk transcriptomes through digital cytometry. This provided a strategy for extending cellular information derived from single-cell data to a larger cohort with matched molecular and clinical information. Rather than treating estimated cell populations as independent biomarkers, the present framework represents TME composition as a multidimensional patient phenotype and incorporates multi-omics relationships among patients. The seven subtypes displayed substantial differences in the relative abundance of myeloid cells, CAFs, malignant epithelial cells, plasma cells, T cells, and B cells.

Transcriptional analyses provided further biological insight into this organization. Our analysis showed that the two ecological families differed significantly in their epithelial characteristics, but not in their overall cytotoxic immune activity. However, when individual subtypes were examined, significant differences emerged in both epithelial characteristics and cytotoxic immune activity, revealing biological diversity within the broader ecological families.

Genome-wide DE analysis further demonstrated extensive transcriptional differences between the ecological families. A total of 1,535 genes were differentially expressed at FDR < 0.05 and |log2 fold change| ≥ 0.58. Among these, *ANXA3, SDCBP2, MUC1, PRSS8, TM4SF1, ENC1, SPATS2L, TMBIM1*, and *MYO1E* showed higher expression in Family 1 and *ATP2B1-AS1, TRIM52-AS1, PACSIN1*, and *FAM117A* showed higher expression in Family 2. Targeted analyses similarly identified family-associated differences in genes related to epithelial identity, inflammatory and immune signaling, oncogenic signaling, and hypoxia, including *CXCL8, KRT19, KRAS, SMAD4, CXCL10, HIF1A*, and *EPCAM*.

Overall survival differed significantly both between the two ecological families and across the seven individual subtypes. Family 2 showed higher survival status than Family 1, with a log-rank *P* value of 5.7 × 10^−4^, while survival also differed across individual subtypes (*P* = 0.0043). The presence of survival differences at both levels suggests that the broader ecological organization and finer subtype-specific states may each contain clinically relevant information.

Several aspects of our GCN framework distinguish it from conventional feature-based patient stratification. Rather than concatenating heterogeneous variables into a single high-dimensional matrix, separate patient-similarity graphs preserve modality-specific relationships among tumors. Our model was guided toward the TME through both the fusion prior and reconstruction-loss weighting, consistent with the study’s objective of investigating microenvironment-associated PDAC heterogeneity. We are aware that the TME-guided weighting strategy intentionally biases representation learning toward microenvironmental information. However, our sensitivity analyses support a strong contribution of the TME.

Our study has some limitations. First, subtype discovery and downstream biological and survival analyses were performed within TCGA dataset, and independent validation will be necessary to establish the reproducibility and generalizability of the seven-subtype and two-family classifications. Second, TME composition was computationally inferred from bulk transcriptomic data rather than measured directly in the same TCGA tumors at single-cell resolution. Although the reference was derived from PDAC single-cell data, deconvolution necessarily simplifies cellular diversity into a limited number of reference populations. Finally, the identified ecological families and transcriptional programs represent computational associations and will require experimental and prospective clinical validation before they can be considered therapeutically actionable.

Future studies could extend this framework in several directions. Larger independent PDAC cohorts with matched molecular and clinical data could be used to validate subtype reproducibility and evaluate multivariable associations with patient outcome. Higher-resolution single-cell references could further resolve immune, fibroblast, and malignant-cell states rather than representing them as broad cellular compartments. Integration of spatial transcriptomic information could additionally determine whether the ecological states identified computationally correspond to reproducible spatial organizations within pancreatic tumors. Finally, evaluating treatment response across the identified subtypes could determine whether TME-guided patient stratification provides information relevant to therapeutic selection.

In conclusion, our study demonstrates that integrating single-cell-informed TME composition with complementary clinical, epigenomic, and genomic patient-similarity networks through multiplex graph learning can reveal structured heterogeneity within PDAC. The resulting seven patient subtypes capture distinct microenvironmental and transcriptional states and organize into two broader ecological families associated with differences in overall survival. The strong contribution of the TME across model configurations further emphasizes the importance of incorporating tumor ecosystem information into molecular patient stratification. These findings provide a framework for linking high-resolution single-cell information with larger multi-omics cohorts and establish a foundation for future validation of TME-informed PDAC classification in independent and clinically characterized patient populations.

## DATA AVAILABILITY

The multi-omics datasets analyzed in this study, including gene expression, DNA methylation, CNV, and clinical data, were obtained from TCGA through the Genomic Data Commons Data Portal (https://portal.gdc.cancer.gov/) using the R package *TCGAbiolinks* (project accession: TCGA-PAAD). Single-cell RNA sequencing (scRNA-seq) datasets were obtained from the NCBI Gene Expression Omnibus under accession numbers GSE194247 (immune cells) and GSE235449 (non-immune cells). The supplementary materials and source code for our pipeline, including data preprocessing, analysis, and results generation, are available at *https://github.com/shayorib/PDAC_GCN*.

